# funkea: Functional Enrichment Analysis in Python

**DOI:** 10.1101/2024.08.24.609502

**Authors:** Benjamin Tenmann, Karina Close, Andrea Rodriguez-Martinez, Samer Abujudeh

## Abstract

Advancements in Genome-wide association studies (GWAS) have led to the discovery of numerous genetic variants potentially linked to various traits, necessitating effective methods to interpret and summarise these vast data sets. We introduce `funkea`, a Python package designed to fill this need by providing functional enrichment analysis methods. This tool encompasses popular enrichment approaches under a unified interface and leverages Spark for virtually limitless scale. This allows researchers to conduct pathway, cell-type, and tissue enrichment analysis across diverse annotation datasets. Ultimately, the `funkea` Python package delivers a highly flexible and scalable solution for functional enrichment analysis in the context of modern genetics workflows. https://github.com/BenevolentAI/funkea

## 1 Introduction

Genome-wide association studies (GWAS) have proven themselves as an effective way of linking human genetic variation to disease. GWA studies are now being produced at increasingly large scales, surfacing thousands of potentially causal genetic variants for various traits. With this scale comes a need to introspect and summarise GWAS results and to link them back to the underlying biology. One family of methods is functional enrichment analysis, which attempts to surface sets of genomic functional annotations which are likely to be disproportionately affected in some way by the trait-associated variants (enrichment). Many methods have been proposed for this purpose; however, these methods are generally developed as command-line tools and lack the scalability needed to handle modern genetics workflows.

For this purpose we developed `funkea`, a Python package containing popular functional enrichment methods, leveraging Spark for effectively infinite scale. All methods have been unified into a single interface, giving users the ability to easily plug-and-play different enrichment approaches.

## 2 Methods

`funkea` unifies five popular enrichment methods. It does so by identifying that each method consists of

1. A *data pipeline* — a series of steps transforming user input into information usable by the enrichment method. Inputs are always *at least* a space of genomic annotations 𝒢 (and its subsets 𝒮_*i*_ ∈) 𝒢 and the summary statistics of at least one GWAS.
2. An *enrichment method* — a method consuming the outputs of the data pipeline and returning enrichment results and the significance thereof, for each 𝒮_*i*_.

These two steps are wrapped by a *workflow*, which manages the execution. For demonstration, we analysed the enrichment results of the different methods for the OpenGWAS Parkinson’s study (ID: `ieu-b-7`; figure 1) and a cholesterol study using UK Biobank data.

**Table 1:** The available enrichment methods and the partition types for which they work.

| Name | Partition types | Source |
| --- | --- | --- |
| DEPICT | Soft / Hard | Pers et al. [2015] |
| Fisher | Hard | Fisher [1922] |
| GARFIELD | Hard | Iotchkova et al. [2019] |
| S-LDSC | Hard | Finucane et al. [2015] |
| SNPsea | Soft / Hard | Hu et al. [2011] |

**Figure 1:**
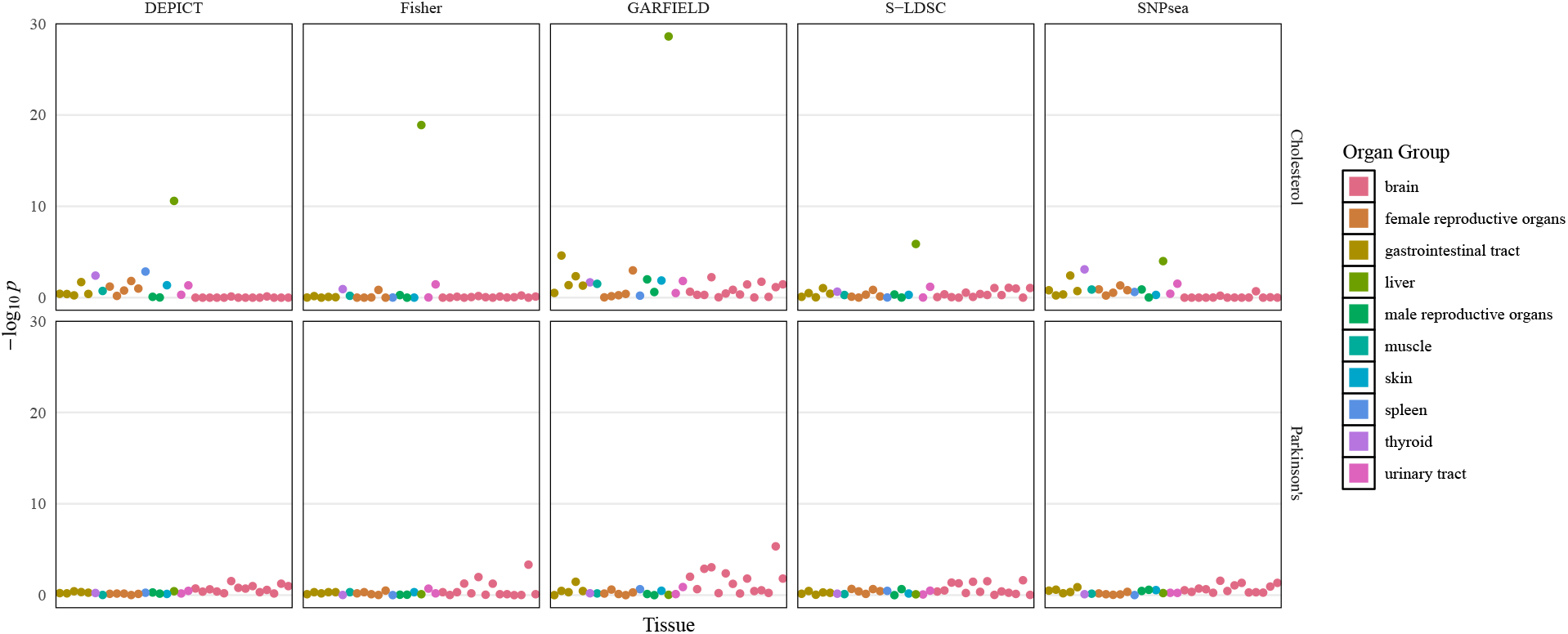
Log-transformed *p*-values of the tissue enrichments from GWA studies on cholesterol and Parkinson’s disease. The annotation data used here was a subset of GTEx [Lonsdale et al., 2013].

### 2.1 Data Pipeline

Each method has its own data pipeline, taking at least an annotation component and the summary statistics of a GWA study. The annotation component defines the space of genomic annotations 𝒢, which is a set sequence spans with coordinates defined genome-wide; that is, we know their start and end base-pairs, and chromosome. A classic example of genomic annotation is the gene, but in general, any sequence span is valid. These annotations are expected to be partitioned into *K* subsets 𝒮_*i*_ ∈ 𝒢, ∀*i* ∈ {1, … , *K*} , for which we will get the final enrichments. This could be any biologically relevant partition of the annotations (e.g. pathways, cell-types, tissues etc.), and need not be non-overlapping. Also, for some methods partitions can be ‘soft’, i.e. a given annotation can have a distribution over the partitions. For example, gene expression values for different tissues can be viewed as unnormalised probabilities.

### 2.2 Enrichment Method

Once the data have been wrangled into the appropriate input format, the enrichment methods compute the enrichment and the corresponding significance for each partition *i* of the annotations. Enrichment is uniquely defined for each method (overview in appendix). Equally, significance is established differently, as the null hypothesis varies from method to method. However, to allow for plug-and-play, each method returns a single dataframe, containing the enrichment value and corresponding *p*-value for each partition *i*.

### 2.3 Additional Features

To be able to run all of these methods at scale, some additional tooling needed to be implemented. Users may find these useful:

1. **LD pruning** — A Spark UDAF written in Scala to run LD pruning seamlessly at any scale. Required in: `DEPICT`, `GARFIELD`
2. **Hypergeometric test** — A Spark implementation of the hypergeomtric test. Required in: `Fisher`
3. **(stratified) LD score regression** — A Python 3 implementation of stratified LD score regression, compatible with Spark. Required in: `S-LDSC`

## 3 Discussion

In conclusion, the `funkea` Python package gives users an easy way to run functional enrichment analysis at any scale. Each enrichment method can be applied flexibly to any annotation dataset, allowing users to run pathway, cell-type and tissue enrichment analysis with the same set of tools.

## 4 Acknowledgments

This research has been conducted using the UK Biobank Resource under Application Number 43138. Using real patient data is crucial for clinical research and to find the right treatment for the right patient. We would like to thank all participants who are part of the UK Biobank, who volunteered to give their primary and secondary care and genotyping data for the purpose of research. UK Biobank is generously supported by its founding funders the Wellcome Trust and UK Medical Research Council, as well as the British Heart Foundation, Cancer Research UK, Department of Health, Northwest Regional Development Agency and Scottish Government.

## A Appendix

### A.1 Fisher’s exact test

The hypergeomtric test, or Fisher’s exact test [Fisher, 1922], is a naïve functional enrichment method, which computes the significance in the overlap between ‘enriched’ subset 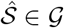 and ‘true’ subsets 𝒮_*i*_ ∈ 𝒢, ∀ *i* ∈ {1, … , *K*}. Hence, the enrichment *e*_*i*_ of a given study for partition *i* of 𝒢 is defined as

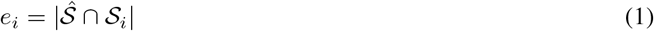

where 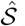 is the set of *s*_*ij*_ overlapped by all the genome-wide significant variants in the study. The significance of *e*_*i*_ is assumed to be distributed hypergeomtrically, and hence

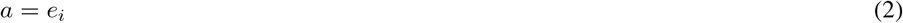

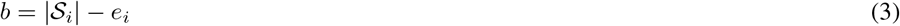

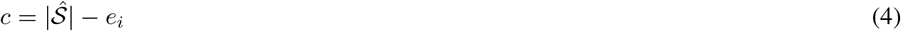

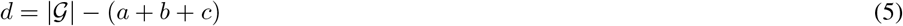

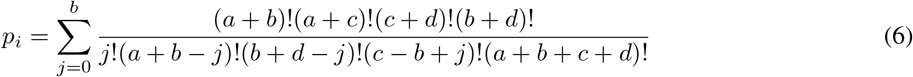

The Fisher’s method can be extended with linkage disequilibrium (LD) pruning (for variant selection) and / or FDR correction (for enrichment selection). However, we have found these modifications generally did not improve enrichment results.

### A.2 SNPsea

Originally developed for tissue and cell-type enrichment [Hu et al., 2011, Slowikowski et al., 2014], `SNPsea` has also been applied to pathway enrichment [Slowikowski et al., 2014]. It uses activity matrix **X** to allow for definitions of ‘soft’ partitions; that is, distributing each annotation over the *K* partitions. For example, in tissue enrichment **X** may be a gene expression matrix such that *X*_*ij*_ is the expression of gene *j* in tissue *i*. A nice property of this method, is that it is defined both for **X** ∈ ℝ^*K×*|*G*|^ and **X** ∈ {0, 1} ^*K×*|*G*|^, i.e. it works for soft *and* hard partitions. If **X** ∈ ℝ^*K×*|𝒢|^, then it is normalised in the following way:

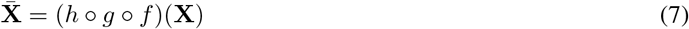

where

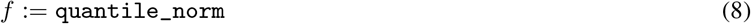

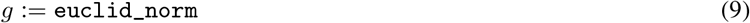

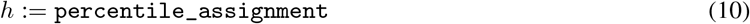

(9) is the easiest to explain mathematically, and simply normalises each column vector in **X** to unit length; that is, each annotation *j* will have length 1 over all *K* partitions. In the example of gene expression, this amplifies specifically expressed genes and surpresses ubiquitously expressed ones.

To better understand (8), consider the following code snippet:

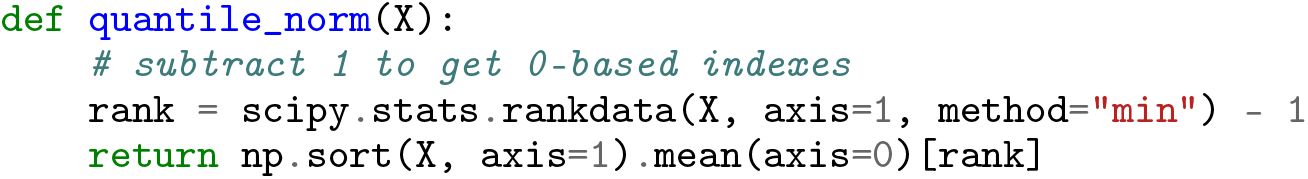

i.e. we average the *observed* quantiles in each partition over the *K* partitions and then assign each annotation in each partition to a quantile. See Amaratunga and Cabrera [2001], Bolstad et al. [2003] for a reference.

Similarly, (10) assigns 1 minus the percentile of each annotation in a given partition to that annotation, such that the final activity matrix 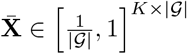. It is important to emphasise that this means that *higher* values in **X** will be assigned to *lower* values in 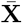.

Next, `SNPsea` defines a locus as the sequence span covered by the furthest LD proxies of a genome-wide significant variant. These loci are then linked to annotations by overlapping the locus spans with the annotation spans. If a locus does not overlap any annotations, it will be expanded by 10kb on either side and overlap will be attempted again. Loci which overlap the same annotations are combined into a single locus.

These annotation-locus links can be represented in a bipartite adjacency matrix **L** ∈ {0, 1} ^|𝒢|*×M*^ , where *M* specifies the number of loci in the study. Using this and 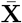, we can compute the partition-locus specificity score matrix **K**, where *K*_*il*_ between partition *i* and locus *l* is defines as:

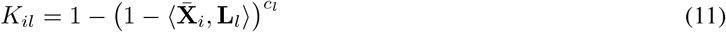

where 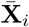 is the normalised activity profile for partition *i*, **L**_*l*_ is the adjacency vector for locus *l* and

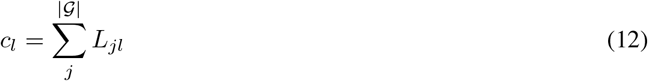

is the total number of annotations in locus *l*. Finally, the inner product between **X**_*i*_ and **L**_*l*_ is defined as:

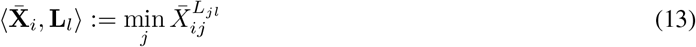

Since *L*_*kl*_ ∈ {0, 1} and 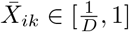, the above inner product will always return the highest specificity value in a locus. Remember that a low value in 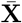 (percentile), denotes high specificity. The enrichment *e*_*i*_ for partition *i* is defined as:

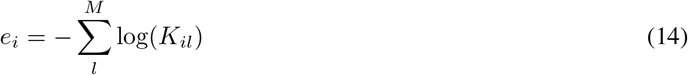

In the case when **X** ∈ {0, 1}^|𝒢|*×M*^ , the specificity score matrix **K** is computed in the following way:

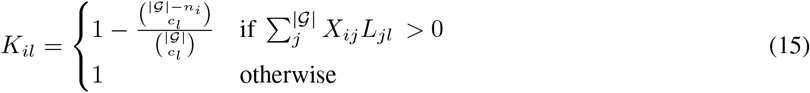

where 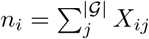.

The significance of *e*_*i*_ is then computed via a permutation-based test. First, a set of ‘null’ loci are generated from a list of LD pruned, non-associated variants. Then, a set of null loci are sampled, such that each true locus has a null counterpart, which contains (roughly) the same number of annotations (i.e.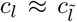). From these null loci, a null enrichment 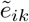 is computed. These steps are repeated *N* times, giving us the *p*-value for enrichment *e*_*i*_, like so:

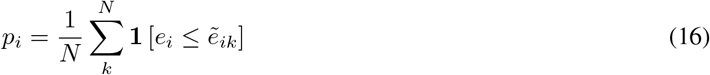

### A.3 GARFIELD

`GARFIELD` is a general method for functional enrichment analysis [Iotchkova et al., 2019]. It regresses locus association to a given trait on whether the lead variant — or one of its LD proxies — overlaps a given partition *i*, as well as a set of controlling covariates. Formally:

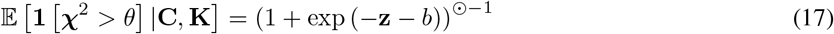

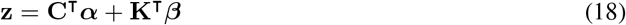

where 𝒳^2^ ∈ ℝ^*M*^ are the association statistics for the LD pruned lead variants, *θ* is some threshold, **C** ∈ {0, 1} ^*D×M*^ are the controlling covariates and **K** is:

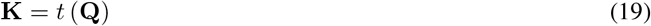

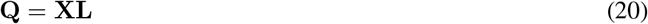

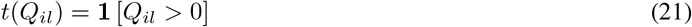

an indicator matrix showing whether a given locus overlaps an annotation partition. The definitions of **X** and **L** are the same here as for `SNPsea`; however, **L** is constructed in a slightly different way. As mentioned above, an annotation *j* and a locus *l* are considered ‘linked’, if the lead variant or one of its LD proxies overlaps it. If the overlap is with an LD proxy, the annotation as to also be withing 500kb of the lead variant.

Here, we define the controlling covariates **C** as being binary; however, any type of variant-level covariate is valid. In the original formulation of `GARFIELD`, **C** is divided into two feature type subsets: (1) binned distance from transcription start site (TSS), and (2) binned number of LD proxies. Formulated differently, **C** is an indicator matrix, telling us how far from the nearest gene a given lead variant is, and how much LD it is experiencing. The enrichment *e*_*i*_ for given partition *i* is then defined as follows:

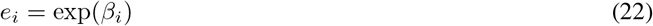

The significance of the enrichment is then computed in the standard way for linear models:

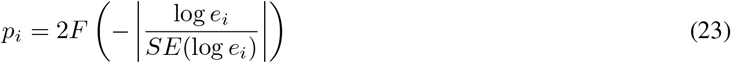

where *F* is the cumulative density function of a unit Gaussian.

### A.4 S-LDSC

Stratified LD score regression has been used for functional and tissue enrichment analysis [Finucane et al., 2015, 2018]. It is defined as follows:

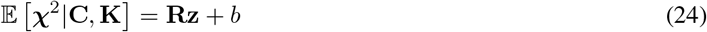

where **R** is the square LD correlation matrix and **z** is defined in (18). Similarly to `GARFIELD`, the enrichment of partition *i* is defined as:

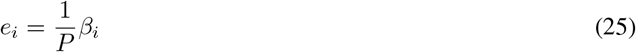

where *P* is the sample size (i.e. number of patients) used in the GWAS. From LD score regression, we can see that this is the total heritability of partition *i* [Finucane et al., 2015]. *From this enrichment we get the p*-value in the usual way:

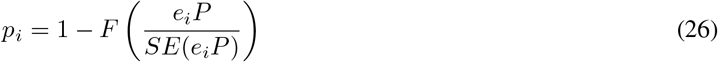

The adjacency matrix **L** is again constructed in a slightly different way. Here, every variant is considered individually, and considered ‘linked’ to a particular annotation if it is within 100kb of that annotation. Moreover, **C** is usually another **K**, derived from a different **X** and **L** — i.e. different annotations and corresponding partitions.

### A.5 DEPICT

`DEPICT` is a popular method for gene prioritisation, but also has proven itself useful for functional enrichment analysis [Pers et al., 2015]. Similarly to `SNPsea`, it uses activity matrix **X** ∈ ℝ^*K×*|𝒢|^ to assign the annotations *s*_*ij*_ to the (soft) partitions *i* ∈ {1, … , *K*} , by normalising the activities over the *K* partitions. However, in this case, 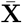 is produced from **X** via *z*-scoring, rather than normalising to unit length. Also, much like `SNPsea`, `DEPICT` defines bipartite adjacency matrix **L** ∈ {0, 1} ^|𝒢|*×M*^ , which defines whether an annotation and a locus overlap. It then computes the scores between each partition *i* and locus as follows:

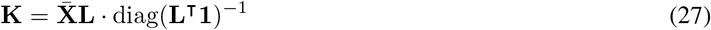

The loci are defined as the LD pruned genome-wide significant variants and their LD proxies. Each locus is overlapped with the genomic annotations and loci which do not overlap any annotations are associated with their nearest annotation. Loci which share annotations are fused.

From **K**, we then get the biased estimate of the enrichment 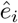 of partition *i*:

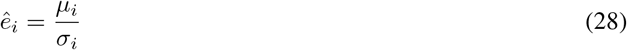

where *µ*_*i*_ and *σ*_*i*_ are the sample mean and standard deviation of the partition-locus scores *K*_*ij*_. The final bias corrected enrichment *e*_*i*_ is then defined as

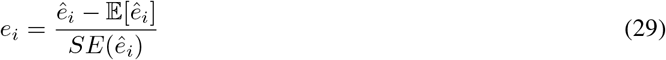

where 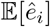 and 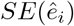 are the expectation and standard error of 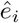, which are approximated empirically via Monte Carlo sampling. Since we assume a normal distribution of *e*_*i*_, we get *p*_*i*_ = 1 − *F* (*e*_*i*_), where *F* (*e*_*i*_) is the cumulative distribution function of the unit Gaussian.

Finally, `DEPICT` estimates the false discovery rate (FDR) for each *p*_*i*_ by running *N* repetitions of the above steps using *M* randomly generated loci. Each generated locus will have the same number of annotations as their ‘real’ counterpart. The FDR is computed as follows:

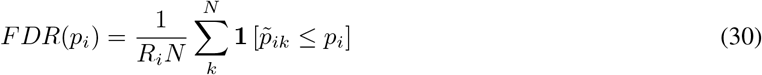

where *R*_*i*_ is the ordinal rank of *p*_*i*_ and 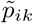 is the *p*-value of the *k*-th null enrichment for partition *i*. For the sake of consistency with the other methods, we did not include this step in our Python package.

## References

Tune H Pers, Juha M Karjalainen, Yingleong Chan, Harm-Jan Westra, Andrew R Wood, Jian Yang, Julian C Lui, Sailaja Vedantam, Stefan Gustafsson, Tonu Esko, et al. Biological interpretation of genome-wide association studies using predicted gene functions. Nature communications, 6(1):5890, 2015.

Ronald A Fisher. On the interpretation of χ 2 from contingency tables, and the calculation of p. Journal of the royal statistical society, 85(1):87–94, 1922.

Valentina Iotchkova, Graham RS Ritchie, Matthias Geihs, Sandro Morganella, Josine L Min, Klaudia Walter, Nicholas John Timpson, UK10K Consortium, Ian Dunham, Ewan Birney, et al. Garfield classifies disease-relevant genomic features through integration of functional annotations with association signals. Nature genetics, 51(2): 343–353, 2019.

Hilary K Finucane, Brendan Bulik-Sullivan, Alexander Gusev, Gosia Trynka, Yakir Reshef, Po-Ru Loh, Verneri Anttila, Han Xu, Chongzhi Zang, Kyle Farh, et al. Partitioning heritability by functional annotation using genome-wide association summary statistics. Nature genetics, 47(11):1228–1235, 2015.

Xinli Hu, Hyun Kim, Eli Stahl, Robert Plenge, Mark Daly, and Soumya Raychaudhuri. Integrating autoimmune risk loci with gene-expression data identifies specific pathogenic immune cell subsets. The American Journal of Human Genetics, 89(4):496–506, 2011.

John Lonsdale, Jeffrey Thomas, Mike Salvatore, Rebecca Phillips, Edmund Lo, Saboor Shad, Richard Hasz, Gary Walters, Fernando Garcia, Nancy Young, et al. The genotype-tissue expression (gtex) project. Nature genetics, 45(6):580–585, 2013.

Kamil Slowikowski, Xinli Hu, and Soumya Raychaudhuri. Snpsea: an algorithm to identify cell types, tissues and pathways affected by risk loci. Bioinformatics, 30(17):2496–2497, 2014.

Dhammika Amaratunga and Javier Cabrera. Analysis of data from viral dna microchips. Journal of the American Statistical Association, 96(456):1161–1170, 2001.

Benjamin M Bolstad, Rafael A Irizarry, Magnus Åstrand, and Terence P. Speed. A comparison of normalization methods for high density oligonucleotide array data based on variance and bias. Bioinformatics, 19(2):185–193, 2003.

Hilary K Finucane, Yakir A Reshef, Verneri Anttila, Kamil Slowikowski, Alexander Gusev, Andrea Byrnes, Steven Gazal, Po-Ru Loh, Caleb Lareau, Noam Shoresh, et al. Heritability enrichment of specifically expressed genes identifies disease-relevant tissues and cell types. Nature genetics, 50(4):621–629, 2018.

